# SleepEEGpy: a Python-based software “wrapper” package to organize preprocessing, analysis, and visualization of sleep EEG data

**DOI:** 10.1101/2023.12.17.572046

**Authors:** G. Belonosov, R Falach, J.F. Schmidig, M. Aderka, V. Zhelezniakov, R. Shani-Hershkovich, E. Bar, Y Nir

**Author notes:** These authors contributed equally to this work (G. Belonosov, R Falach).

## Abstract

Sleep research uses electroencephalography (EEG) to infer brain activity in health and disease. Beyond standard sleep scoring, there is increased interest in advanced EEG analysis that require extensive preprocessing to improve the signal-to-noise ratio, and dedicated analysis algorithms. While many EEG software packages exist, sleep research has specific needs that require dedicated tools (e.g. particular artifacts, event detection). Currently, sleep investigators use different libraries for specific tasks in a ‘fragmented’ configuration that is inefficient, prone to errors, and requires the learning of multiple software environments. This leads to a high initial complexity, creating a crucial barrier for beginners in sleep research. Here, we present SleepEEGpy, an open-source Python software “wrapper” package to facilitate sleep EEG data preprocessing and analysis. SleepEEGpy builds upon MNE-Python, YASA, and SpecParam tools to provide an all-in-one, beginner-friendly package for comprehensive yet straightforward sleep EEG research including (i) cleaning, (ii) independent component analysis, (iii) detection of sleep events, (iv) analysis of spectral features, and associated visualization tools. A dashboard visualization tool provides an overview to evaluate data and its preprocessing, which can be useful as an initial step prior to detailed analysis. We demonstrate the SleepEEGpy pipeline and its functionalities by applying it to overnight high-density EEG data in healthy participants, revealing multiple characteristic activity signatures typical of each vigilance state. These include alpha oscillations in wakefulness, sleep spindle and slow wave activities in NREM sleep, and theta activity in REM sleep. We hope that this software will be embraced and further developed by the sleep research community, and constitute a useful entry point tool for beginners in sleep EEG research.

## Introduction

Electroencephalography (EEG) is the main tool in basic and clinically oriented sleep research (1). EEG is routinely used in conjunction with electrooculography and electromyography to perform sleep scoring and distinguish between vigilance states of wakefulness, rapid eye movement (REM) sleep, and non-REM sleep (2). Sleep scoring is performed either manually according to established standards (3) or, in recent years, via automatic tools (4–6). Beyond sleep scoring, there is increased attention toward advanced EEG analysis that focuses on investigating events occurring at specific times, frequencies, and scalp locations or in estimated brain sources (7). Examples of such sleep EEG analyses include an association between sleep spindles and sleep-dependent memory consolidation (8–13), regional differences in slow-wave activity during development (14), changes in slow-wave-spindle coupling in old age (15), and neural correlates of dreaming (16). In the clinic, advanced analysis that goes beyond sleep architecture reveals an association between disrupted frontal slow waves and β-amyloid pathology in Alzheimer’s disease (17), altered central sleep spindles in schizophrenia (18), and how epileptic seizures emerge from sleep oscillations (19).

EEG data comprise a mixture of a signal of interest from neuronal origin and noise arising from both physiological and extrinsic origins (20). Typically, preprocessing (e.g., filtering and artifact rejection) is performed to improve the signal-to-noise ratio (SNR) of EEG signals (21). While EEG may be “better left alone” in event-related contexts where noise can be reduced by trial averaging (22), it is crucial to preprocess and clean the EEG signal when analyzing the continuous, spontaneous brain activity observed during sleep. For example, many artifacts, such as sweating or eye movements, are dominated by spectral frequencies that overlap with slow-wave activity; hence, removing such artifacts is often necessary to accurately characterize sleep homeostasis as indexed by slow-wave activity (23).

Various established software packages exist for visualization, preprocessing, and analysis of EEG data (24). Leading general-purpose EEG software packages include open-source MATLAB-based Brainstorm (25), EEGLAB (26) and FieldTrip (27), and Python-based MNE-Python (28), all of which are continuously developed and are witnessing a rapidly expanding interest in the scientific community (fig. 1a). EEGLAB and Brainstorm, developed earlier, provide a graphical user interface, whereas Fieldtrip and MNE-Python are scripting oriented. Over the last decade, automatic preprocessing pipelines have been developed and are being increasingly used for rejecting artifacts, problematic time intervals, and “bad” electrodes alongside conventional visual annotation (29–38) (Fig 1b).

**Figure 1.**
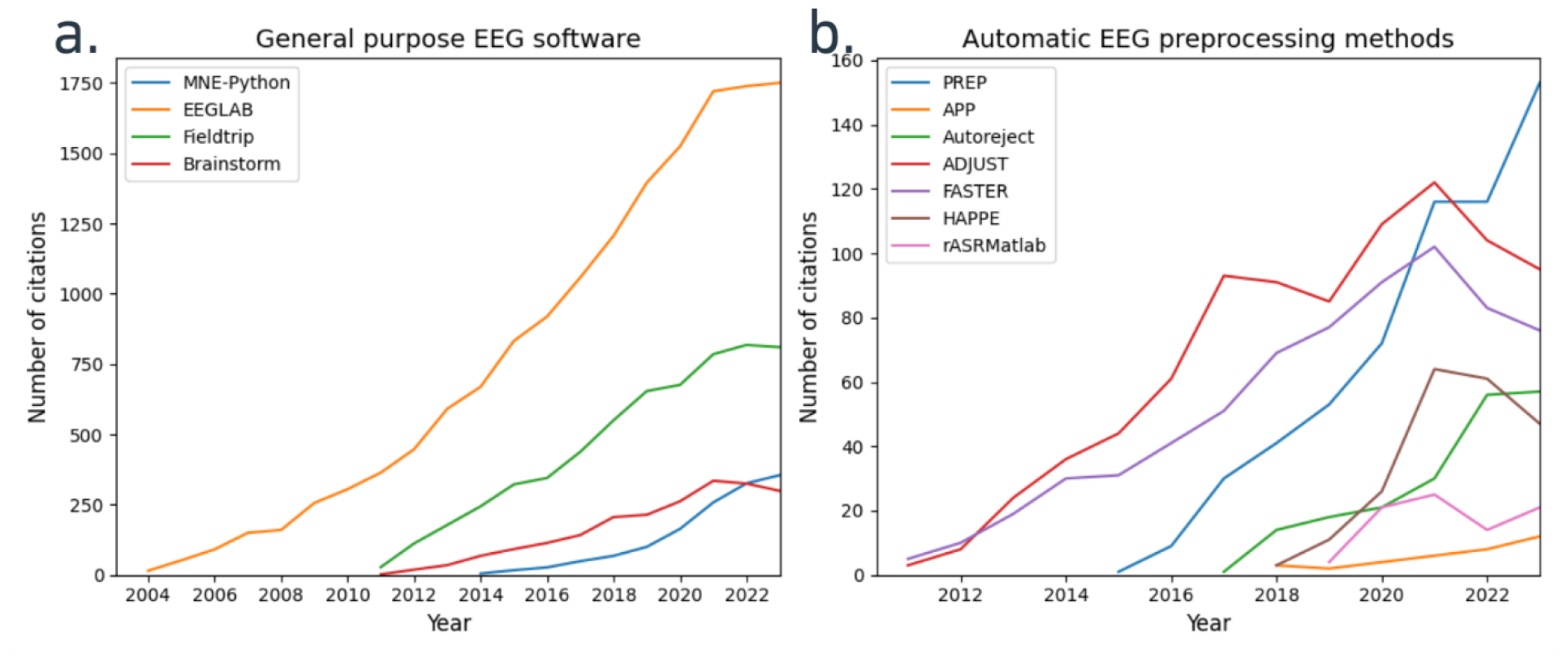
Citation dynamics of common general-purpose and automatic preprocessing EEG software. A noncumulative number of citations (y-axis) per year (x-axis) for (a) four leading general-purpose EEG packages (EEGLAB, orange; Fieldtrip, green; Brainstorm, red; MNE-Python, blue) and (b) leading software packages implementing automatic preprocessing of EEG data (PREP pipeline, blue; APP, orange; Autoreject, green; ADJUST, red; FASTER, purple; HAPPE, brown; rASRMatlab, pink). Citations are based on the Scopus database (17.12.2023).

Sleep EEG research has many specific preprocessing, analysis, and visualization considerations compared with classic event-related potential (ERP) and resting-state EEG studies. For example, different intrinsic artifacts may be dominant (e.g., cardiac activity, muscle twitches, and slow/rapid eye movements), whereas other common artifacts during wakefulness, such as eye blinks, are mostly absent (39). Moreover, sleep EEG activity is mostly spontaneous and not necessarily arranged around specific experimental events, as in ERP studies. Therefore, continuous data are usually not windowed into epochs. This approach warrants preprocessing and visualization that is closer to the raw data, in line with a recently proposed ‘lossless’ preprocessing approach (40). Additional considerations unique to sleep EEG analysis involve sleep stage-based analysis. This includes the detection of specific events such as slow waves and spindles, as well as spectral aspects and their associated scalp topography, which are of interest to most studies employing advanced analysis of sleep EEG. Thus, specific features of sleep EEG present a need for specific software tools tailored to these specific needs. Moreover, the quantity and complexity of these steps can often be overwhelming for beginners to the field.

Ideally, a complete sleep EEG software package should draw on the advantages of both general-purpose tools (e.g., preprocessing, time-frequency, and topography analyses) and specialized tools for sleep research (e.g., integration with sleep scoring, advanced analyses per sleep stage, detection of specific sleep oscillations and other events). At present, this is typically achieved by combining different available and custom-made tools. This “fragmented” configuration is inefficient, prone to errors, and requires the burdensome learning of multiple software environments. These issues are an elevated entry hurdle for new users and often lead to frustrating first encounters with the analysis of sleep EEG data.

Here, we present the SleepEEGpy, a general-case sleep EEG software “wrapper” package that organizes preprocessing, analysis, and visualization of sleep EEG data. We initially developed SleepEEGpy as tool for new students in our lab for working with EEG data of human sleep. It is meant to offer a user-friendly introduction to sleep data analysis for users with little to no prior experience with EEG, sleep or programming. Geared for beginners, it is not meant to replace the rich and complex functionalities of highly developed packages that it is based on (e.g. MNE); rather it facilitates entry point for newcomers in sleep EEG research. At the same time, its standardized visualization allows more experienced users to quickly assess the quality of the applied sleep-scoring, pre-processing and analysis of the sleep data and thus enables them to provide helpful feedback and effectively mentor and supervise new users. SleepEEGpy is based on the following Python packages: MNE(28), YASA (41), and SpecParam (42) (formerly FOOOF, for analysis of periodic and aperiodic activities). The choice of Python as an open-source programming language leverages the benefits of its many libraries, including extended machine learning ecosystems, broad documentation, and a dynamic community. To facilitate rapid learning, SleepEEGpy uses Jupyter notebooks (43). Notably, SleepEEGpy is not meant to replace or improve the packages it is built on. Its value primarily lies in simplifying getting started with these packages by unifying them into one framework and by reducing the functionality to the core necessities for the analysis of general sleep EEG data.

## Methods

### Overview

The SleepEEGpy pipeline is divided into two sections: preprocessing and analysis. Section A (preprocessing) is further divided into A1 (cleaning) and A2 (independent component analysis, ICA), whereas section B (analysis) is further divided into B1 (events) and B2 (spectral). The preprocessing section offers to increase the SNR by cleaning (e.g., filtering, rejection of bad electrodes, or problematic temporal intervals) and by regressing out noise components (through an ICA). The analysis section focuses either on specific sleep “events” (graphoelements such as sleep spindles, slow-waves, or rapid eye movements) or power spectral decomposition performed for individual recordings or multiple datasets. The dashboard and additional visualization tools allow a precise and sleep-tailored visual assessment of the preprocessing and the analysis section. Together, SleepEEGpy offers an integrated pipeline for cleaning, ICA, event detection and analysis, spectral analysis, and their visualization, as well as integration with sleep scoring vectors (performed either a priori manually or automatically by linking with YASA).

### Prerequisites: input data, software, and hardware

To utilize SleepEEGpy, input data must consist of non-segmented (‘continuous’) EEG data in any common format supported by MNE-Python, e.g., Brain Vision, Meta File Format (MFF), or European Data Format (EDF). For sleep-stage-based functionality (both events and spectral), an additional sleep scoring vector is required in the form of a text file containing an integer per row representing different sleep stages for each epoch. For example, the sleep module of the Visbrain package (64) provides an interface for sleep-scoring that is well suited and compatible with SleepEEGpy. If not provided, SleepEEGpy can perform automatic sleep scoring based on YASA. We highly recommend that users install SleepEEGpy into Python’s virtual environment (and not the global one) through venv (Python’s built-in), conda environments, or their easier-to-use wrappers in Visual Studio Code or PyCharm. Particularly for long overnight (6-10h) high-density (128/256-channel) EEG sleep datasets, we highly recommend at least 64 GB of rapid access memory (RAM), especially for event detection tools, even when the sampling rate is not higher than 256 Hz.

### Architecture and typical workflow

Each tool within the pipeline is organized independently and has a corresponding Jupyter notebook. These notebooks serve as exemplars, providing a detailed walkthrough of each tool’s functionality and offering step-by-step guidance to new users. More experienced users can always re-organize, reuse, and combine different pipeline tools to support their needs. Figure 2 depicts a possible prototypic process flow of the SleepEEGpy pipeline.

**Figure 2.**
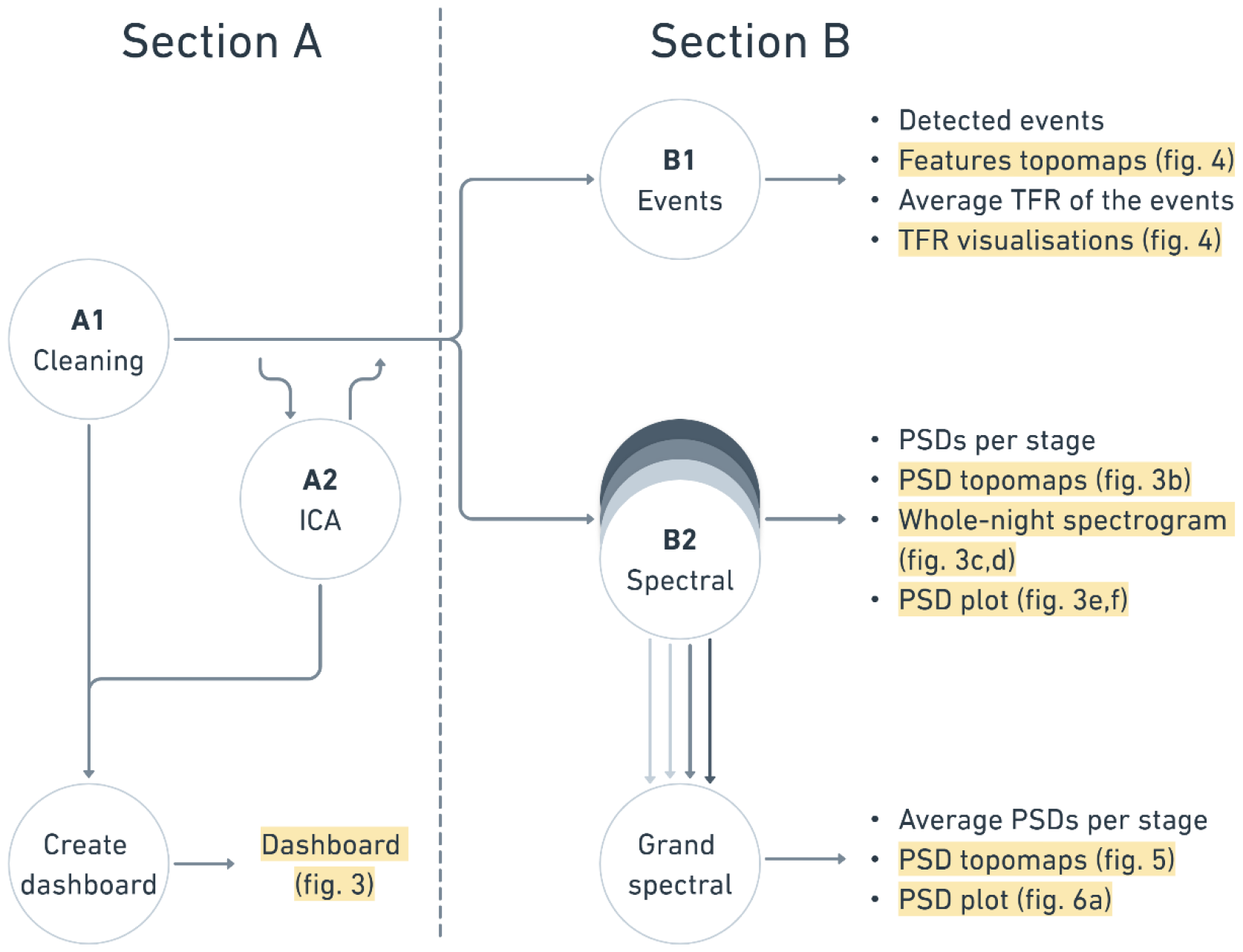
General structure of the SleepEEGpy pipeline. Section A (preprocessing) includes A1, a cleaning tool that takes EEG data as input and (i) performs basic operations such as resampling and filtering according to default or user-defined parameters. (ii) It allows the user to visualize and annotate bad electrodes or bad temporal intervals to refrain from their subsequent analysis. Preprocessing can also include A2, a tool for independent component analysis (ICA) visualizations and component rejection. A ‘dashboard’ (fig. 3) provides a visual summary of preprocessing. Section B (analysis) includes B1, a tool for the detection of specific sleep events (e.g., sleep spindles, slow waves) and their properties, and B2, power spectrum analysis for a single recording and multiple datasets (‘Grand spectral’); overlapping closed discs and multiple arrows illustrate the transition from single recording to multiple dataset analysis. Arrows represent transitions between different pipeline tools. Text blocks represent the main outputs (visualizations are highlighted). PSD, power spectral density; TFR, time frequency representation.

### Section A: Preprocessing

The preprocessing section (Fig 2A) is divided into two tools: A1, cleaning, and A2, ICA. The dashboard offers a convenient way of reviewing the quality of the applied preprocessing steps. Furthermore, we provide a Jupyter notebook for the cleaning (A1) that can be applied with minimal knowledge of sleep and without further manual inputs. It uses default parameters for resampling, filtering and automatic sleep-scoring. Naturally, default parameters will not result in ideal preprocessing, which has to be evaluated carefully. However, it provides a first-pass “low-entry” point for beginners to start exploring preprocessing of sleep EEG and refine further via additional iterations.

### A1. Cleaning

The pipeline cleaning tool consists of the following signal processing techniques: EEG resampling, filtering, re-referencing, noisy channels and temporal intervals annotation and interpolation of the annotated channels. Resampling often is a necessary step in sleep EEG preprocessing because overnight recordings may require tens of gigabytes of RAM. Filters and data annotations provide a way to improve the SNR by removing artifacts from the signal. For example, by applying a notch filter to remove electrical line noise (50/60Hz) or by annotating the intervals of movements to avoid their subsequent analysis. Default parameters, along with their rationale, are presented below. Re-referencing EEG data to the common average is typically used for assessing topography distributions after interpolating noisy channels to ensure, among other things, that the mean is not dominated by outliers. By default, SleepEEGpy provides an MNE-based manual annotation data browser, which we use for visual annotation of noisy electrodes and temporal intervals. However, users can also easily access the MNE object and annotate data programmatically.

### A2. ICA

The ICA tool is MNE-based and consists of EEG signal decomposition, selection of artificial components based on a data browser and components’ topographies, and subsequent EEG signal reconstruction after regressing out the artificial components. EEG, and sleep EEG specifically, contains noise sources unrelated to brain activity. We can identify, and to some extent separate, the noise from the neuronal signal. For example, physiological noise (e.g., electrocardiograph, sweating, rolling eye movements) or external noise (e.g., 50/60Hz line noise) can often be reliably identified based on the components’ topographies and time series. Applying ICA is optional because there are advantages and disadvantages in removing or keeping some components, depending on the subsequent analysis and the focus of the investigation. For example, eye movement-related potentials during sleep may mask neuronal activities and be chosen to be removed in some contexts; however, in other studies, the research question may require their detection (e.g., (44)). Most sleep EEG studies do not employ ICA and prefer to discard entire 20s/30s segments based on manual identification given the rich, long datasets. However, a disadvantage of this approach is that it may limit artifact identification to the few channels used for sleep scoring (45).

### Default parameters

By default, the parameters for preprocessing and visualization are set as follows:

- Band-pass filter: the default is high-pass (but not low-pass) filtering of the data with a cutoff of 0.3 Hz. As defined by MNE, the default filter is a zero-phase (non-causal) finite impulse response (FIR) filter using the window method with the hamming window. Stricter high-pass filtering, e.g., 0.75 Hz, may be preferable in the presence of high-amplitude sweat artifacts (45).
- Notch filter: By default, we set 50 Hz and its harmonics to be filtered using a notch filter. Like the band-pass filter, the notch filter is a zero-phase FIR filter with a hamming window and 1 Hz width of the transition band.
- ICA: The default ICA algorithm, based on MNE’s implementation, is set to FastICA (46), but it can be changed to Infomax (47) and Picard (48). The number of largest-variance PCA components passed to the ICA algorithm is set to 30 by default. Following MNE’s recommendation, the signal is by default high-pass filtered at 1.0 Hz before fitting to reduce the ICA algorithm sensitivity to low-frequency drifts.

### Dashboard: summary and visualization of preprocessing

To provide a visual summary and overview of sleep EEG data preprocessing, we created a dashboard (Fig 3). It accepts as an input the output of either A1 or A2 (fig 2A). Hence, the dashboard can visualize cleaned EEG recordings with or without excluded ICA components. Furthermore, one can optionally refine the visualization by adding a sleep-scoring vector. Based on the input, all visualizations in the dashboard are then computed with or without ICA or sleep scoring. As a pre-configured visualization tool, it displays the characteristic sleep properties of the EEG recording, such that the data quality and SNR of that sleep dataset can be estimated relatively quickly from a “bird’s eye” view. The dashboard includes four sections: general preprocessing information (fig. 3a), topographical power distribution of key oscillations in specific vigilance states (fig. 3b), time–frequency decompositions (spectrogram, computed using the Multitaper (49) method) superimposed with the corresponding hypnogram, once before (fig. 3c) and once after (fig. 3d) the rejection of bad time intervals (and after ICA, if used), and power spectral density (PSD) plots before and after the preprocessing (fig. 3e,f). A full description of the dashboard results for representative data of overnight sleep in a healthy subject is detailed below in the Results section.

**Figure 3.**
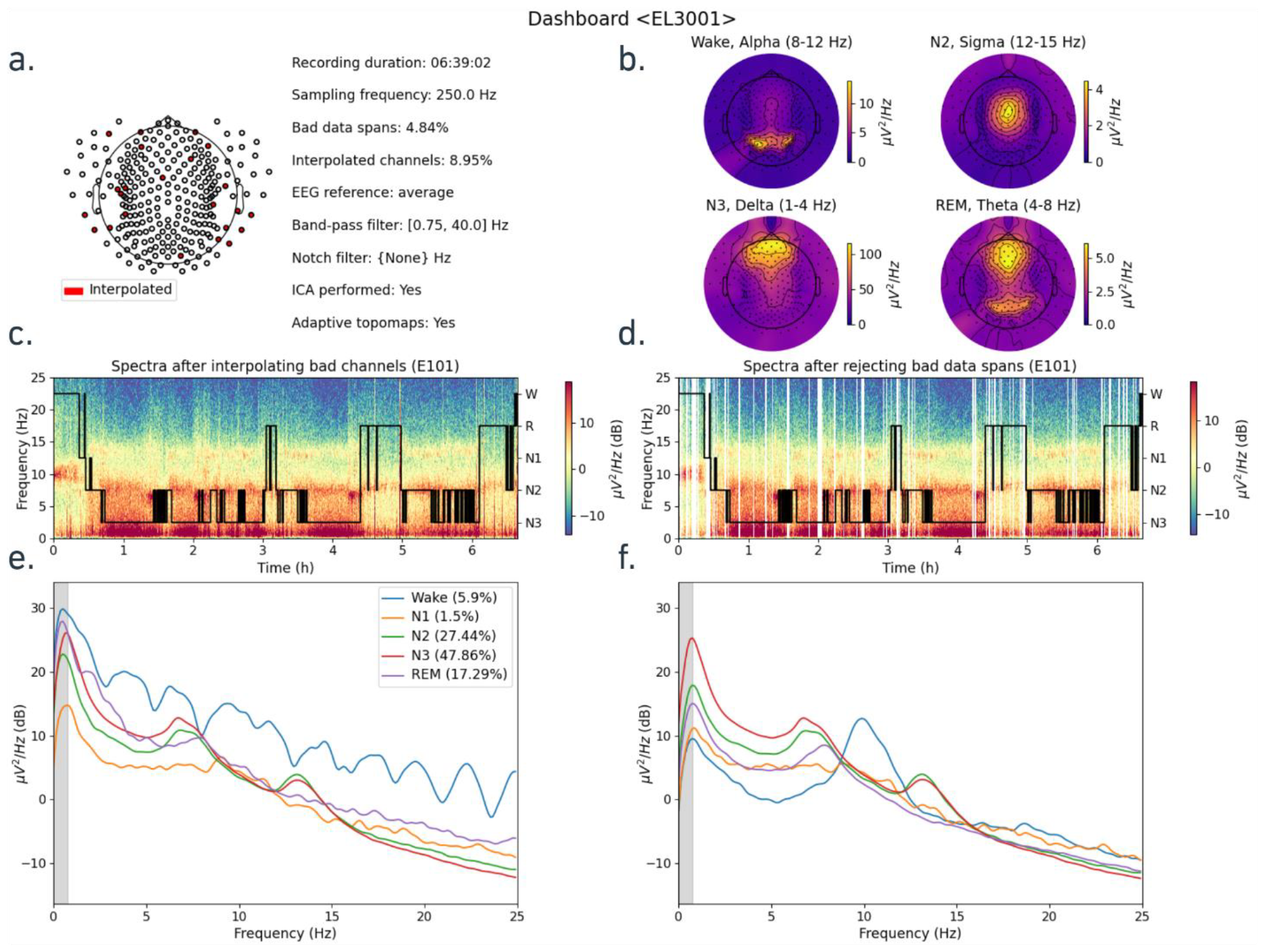
Preprocessing assessment with the dashboard. **a)** left: EEG net montage with spatial distribution of the interpolated channels (in red); right: general information of the recording’s preprocessing. **b)** Topographical PSD distribution per sleep stage in its characteristic frequency band (8-12 Hz for Wake, 12-15 Hz for N2, 1-4 Hz for N3, and 4-8 Hz for REM); subfigures **c–f** are based on the E101 electrode (Pz); subfigures **c)** and **d)** present single-channel spectrogram of the recording (colored power distribution in logarithmic scale over time and frequency) overlapped with a hypnogram (black line representing depth of sleep as a function of time). **c)** spectrogram of the signal after filtering and bad channel interpolation **d)** spectrogram after filtering, bad channel interpolation, bad data span rejection, and optionally, exclusion of artificial independent components (ICA). Subfigures **e)** and **f)** present single-channel power spectral distribution (PSD) plots (as a function of frequency) with lines representing different sleep stages: blue for Wake, orange for N1, green for N2, red for N3, and purple for REM. The gray area on the left of the plots shows frequencies filtered by the high-pass filter. Similar to spectrograms, **e)** is a PSD plot of the signal after filtering and bad channel interpolation, and **f)** is a PSD plot of the signal with additional bad data span rejection and, optionally, exclusion of artificial independent components.

### Section B: Analysis

The analysis section (fig. 2B) is divided into two tools: B1, event-based analysis, and B2, spectrum-based analysis. B1 is mainly based on YASA and B2 is based on MNE and SpecParam. Here we describe the main functionalities, but if fine-tuning of additional parameters is needed it is best to refer to the documentation of the original package. The major advantage here is that SleepEEGpy offers the event- and spectrum-based analysis of sleep EEG within the same framework as the preprocessing and thus is easier to get started for new users. Nevertheless, if one is already acquainted with YASA, employing the package directly might be more convenient than submitting the parameters to YASA through SleepEEGpy.

### B1. Event detection and analysis tools

Event-based tools of the SleepEEGpy pipeline are used to detect and characterize three different sleep graphoelements, namely sleep spindles, slow waves, and rapid eye movements. Each tool takes an MNE-readable EEG file and the sleep scoring vector as input. The input EEG file is typically the output of the preprocessing section, but this is not mandatory. The detection methods of the event-based tools are completely based on YASA detection algorithms (41) and allow the same input parameters. The output of the detection algorithm provides features of the detected events (for example, number of events, amplitude/frequency characteristics of sleep-spindles), as well as the average time–frequency representation averaged per channel across the detected events. The associated visualizations (as seen in fig. 4) include average event time-course plots, topographical distributions of the events’ features, and time–frequency representations.

**Figure 4.**
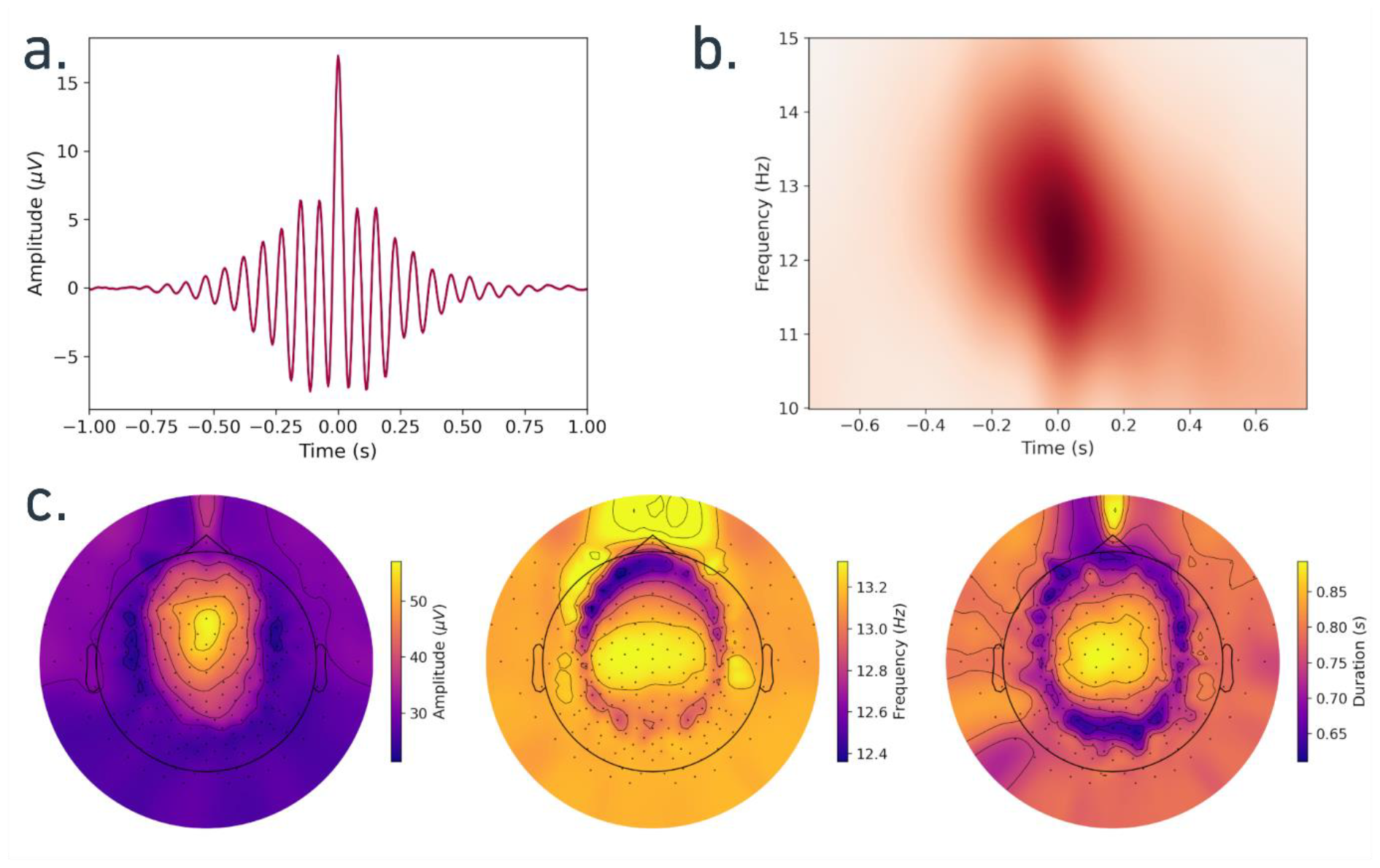
Visualizations of the detected spindles. Overall, 48057 spindles were detected in 257 channels of 193min N2 sleep. (a) Average signal over all detected spindles in all channels, centered around the spindle peak. (b) Average time–frequency representation of detected spindles from channel E101 (Pz). (c) Average topographical distribution of spindle characteristics: amplitude (left), frequency (middle), and duration (right).

### Sleep spindles

are detected using YASA’s algorithm. The algorithm is based on Lacourse et al. (50) and is similar to our previous work (51). In brief, detection is performed by the algorithm using a combination of different thresholds (relative sigma power, root mean square, and correlation) separately for the broadband (by default 1-30 Hz) and sigma-filtered (by default 12-15 Hz) EEG signals. Detection signals are resampled to the original time vector of the EEG data using cubic interpolation to facilitate better precision for spindle start time, end time, and duration. The relative sigma power (relative to the total power in the broadband frequency) is computed using a short-term Fourier transform. Each spindle’s median frequency and absolute power are computed using the Hilbert transform. Additional spindle properties are computed (e.g., symmetry index) as suggested in (52). Detected events with a duration within the range of a prototypical spindle are kept for subsequent analysis (default duration is 0.5-2 s). Finally, an isolation forest algorithm is optionally applied to reject outlier. The EEG signals of the detected spindles can be extracted with a corresponding function, and the properties of the spindles can be accessed through a summary Pandas (53) dataframe.

### Slow waves

are detected via YASA’s algorithm, which is based on a study by Massimini and colleagues (54) and similar to our own previous work (55). In brief, the EEG signal is band-pass filtered in the default range of 0.3–1.5 Hz using a FIR filter with a transition band of 0.2 Hz. Next, negative peaks in the filtered signal with an amplitude of [-40 to -200] uV are detected, as well as all positive peaks with a default amplitude of [10 to 150] uV. For each negative peak (slow wave trough), the nearest positive peak is found, and several metrics are computed, such as the peak-to-peak amplitude, durations of the negative and positive phases, and frequency. A set of logical thresholds, e.g., phase duration, is applied to determine the true slow waves. A Pandas dataframe is created, where each row is a detected slow wave, and each column represents a property of this slow wave. An optional automatic outlier rejection is applied to further remove abnormal slow waves. Similar to the spindle detection algorithm, the EEG signals of the detected slow waves can be extracted.

The detected spindle and slow wave events can be further analyzed using the average time– frequency representation (TFR). Either Morlet wavelets (57) or discrete prolate spheroidal sequence (DPSS) tapers (58) can be used for the TFR computation as implemented by MNE. For the TRF computation of the detected events, we first extracted the EEG signal with a user-defined duration (by default, -1 to 1 sec around the central peak for spindles and negative peak for slow waves). Second, the extracted events are used to compute the average TFR, where the TFR is averaged over events in each channel and each sleep stage. Finally, the average TFR is placed inside an MNE-based container (‘AverageTFR’). It provides additional extensive data manipulation functionality and various visualizations (e.g., fig 4b).

### Rapid eye movements (REMs)

are also detected using YASA’s algorithm. The algorithm uses the rapid eye movement detection method based on left and right outer canthi (left: LOC, right: ROC) EOG data proposed in (56). In brief, the algorithm uses amplitude thresholding of the negative product of the LOC and ROC signals. The REM peaks are detected based on the user-defined frequency range (defaults to 0.5-5 Hz), amplitude threshold (defaults to min 50 and max 325 uV), and REM duration (defaults to min 0.3 and max 1.2 seconds). Similar to spindle and slow wave detection algorithms, the outliers of the detected REMs can be removed using the isolation forest algorithm, the summary Pandas dataframe and their EEG signal can be extracted.

### B2. Spectral tools

Spectral-based tools in the SleepEEGpy pipeline include single recording or multiple dataset analyses. The single recording spectral tool takes an MNE-readable EEG file as input, and the sleep scoring vector (if not provided, can be automatically predicted by the tool with

YASA’s algorithm). The spectral tool for multiple dataset analysis then takes multiple instances of the single recording tool as input. The numeric output of both tools is PSDs calculated separately for each sleep stage. The PSDs per sleep stage are computed using Welch’s method (59) in the following way: first, the EEG signal is divided into regions according to changes in sleep depth, i.e., at the end of the division, there can be multiple segments for each sleep stage. Then, using Welch’s method, PSDs are separately computed for each segment. Finally, separately for each sleep stage, a weighted arithmetic mean based on the length of a segment is applied to the PSDs, producing a per-sleep-stage PSD. The PSD computation is based on the MNE’s Welch function (psd_array_welch), accepts all the original parameters, and uses the same defaults (e.g., the length of FFT and Welch’s segments are set to 256, the segments’ overlap is 0, the default window is the hamming type). In addition, the multiple dataset spectral tool averages PSDs over recordings. Finally, the computed PSDs per sleep stage are placed inside MNE-based containers (‘SpectrumArray’) to preserve the rich spectrum-related functionality of MNE.

PSD results can be visualized either in a “traditional” (power vs. frequency) plot (as in fig. 3e/f or fig. 6a) or with corresponding scalp topography distributions (as in fig. 3b or fig. 5). Additional manipulations and visualizations of the PSDs per sleep stage are available through the MNE containers. In-depth spectral analysis can be conducted with SpecParam (formerly FOOOF) (42) (fig 6b). This analysis improves the characterization of signals of interest by overcoming the limitations of conventional narrowband analyses, e.g., misinterpretation of physiological phenomena (42). This is accomplished by parameterizing neural PSDs into periodic and aperiodic components. This algorithm can identify periodic oscillatory parameters, including the center frequency, power, and bandwidth. In addition, offset and exponent parameters can be extracted for the aperiodic component.

**Figure 5.**
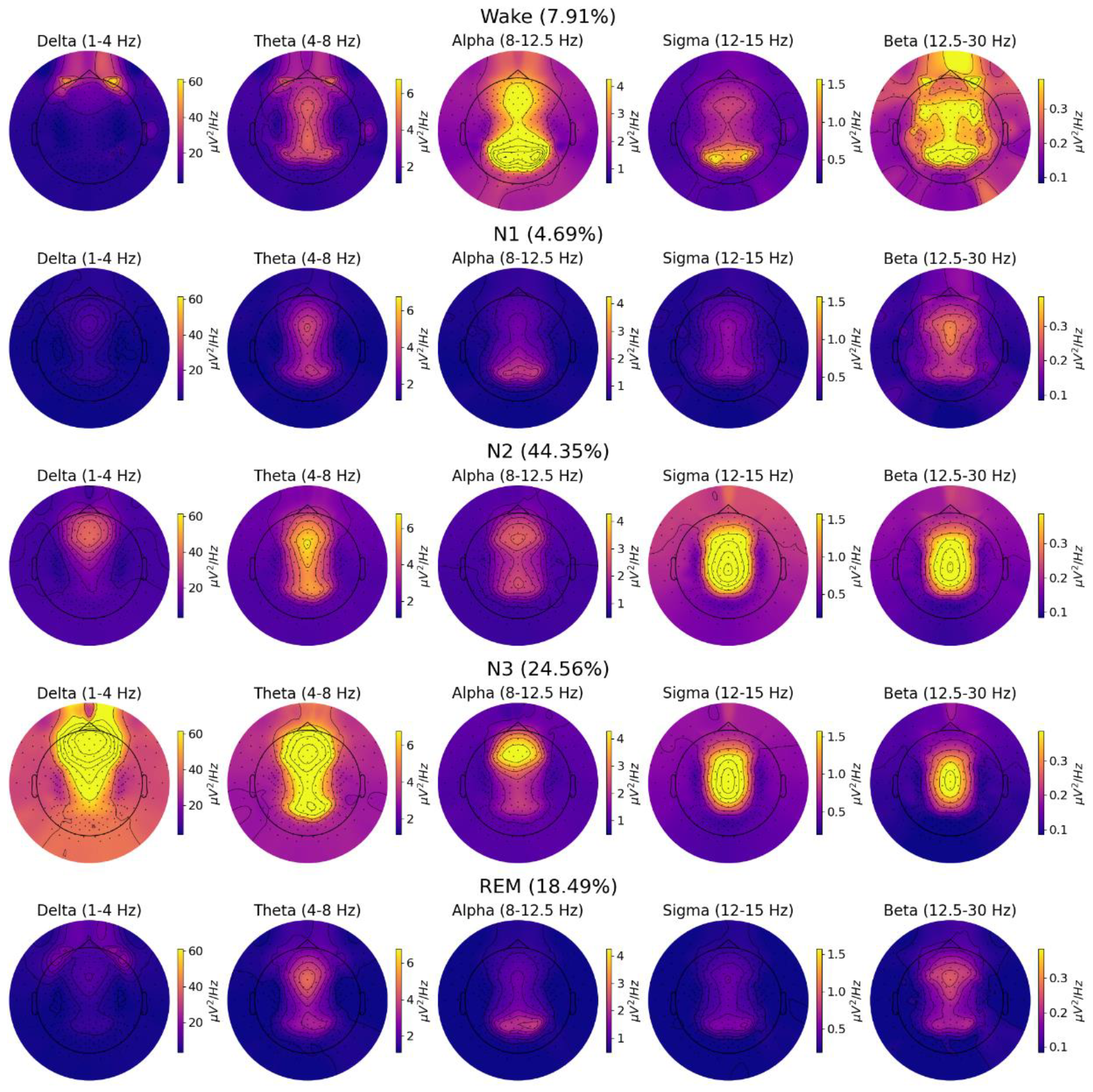
The topographical distribution of PSD per frequency band and sleep stage averaged over 40 subjects. Each row represents a sleep stage, from top to bottom: Wake, N1, N2, N3, and REM. Each column represents a frequency band, from left to right: Delta (1-4 Hz), Theta (4-8 Hz), Alpha (8-12.5 Hz), Sigma (12-15 Hz), Beta (12.5-30 Hz). Percentages in brackets represent a fraction of the sleep stage signal from the overall data.

**Figure 6.**
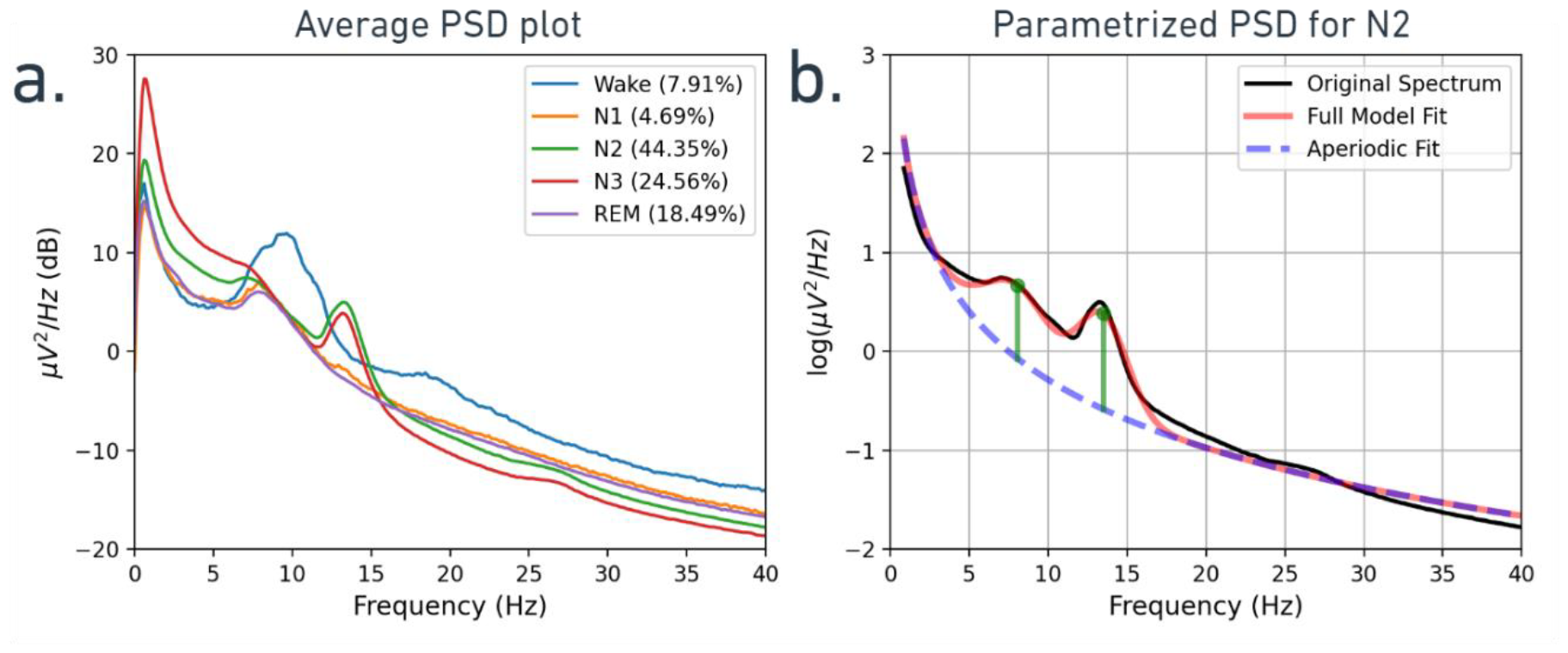
Group-average PSD plots. (a) PSDs from the E101 (Pz) electrode averaged over 40 overnight recordings. The PSDs were transformed to dB for visualization. Percentages in the legend represent the fraction of a sleep stage signal from the overall signal. (b) Average PSD of the N2 stage parametrized with SpecParam. The PSD values were log-transformed. Vertical green lines represent fitted peaks with central frequencies of ∼8.06 and ∼13.47 Hz, and their respective power above the aperiodic component of ∼0.74 and ∼0.97 log(uV^2^/Hz). PSD, power spectral density.

Each spectral tool can be computed and viewed at the single recording level (as in fig. 3e/f) or across multiple datasets (fig. 5,6) recordings, where PSDs are averaged separately across multiple datasets for each EEG electrode and frequency band. Finally, to allow additional preprocessing flexibility and benefit from the interface with diverse approaches (33), spectral tools can also accept an epoch-based signal annotated with sleep stages as an input (as in ERP studies) rather than a ‘continuous’ EEG signal typical for sleep studies.

### EEG data

We illustrate SleepEEGpy functionalities by applying it to sleep recordings of 44 healthy, young adult participants (25 females, age 25.86 ± 3.14 years (mean ± STD), ranging from 21 to 36 years) who participated in a research study on sleep and memory consolidation. Written informed consent was obtained from each participant. This study was approved by the Medical Institutional Review Board of the Tel Aviv Sourasky Medical Center. Participants did not have any history of neuropsychiatric or sleep disorders. Each participant arrived at the sleep lab around 21:00 and, after cognitive testing, proceeded to undisturbed sleep (the data presented here). The mean duration of the recording was 7:18:28 ± 0:36:20 (hours:minutes:sec, mean ± STD). We collected polysomnographic data, including high-density electroencephalogram (hd-EEG), EOG, EMG, and video, as described in (10). hd-EEG was recorded using a 256-channel (plus one reference channel) hydrogel geodesic sensor net (Electrical Geodesics, Inc. [EGI]). Signals were referenced to Cz, amplified using an antialiasing filter and an AC-coupled high input impedance amplifier (NetAmps 300, EGI), and digitized at 1000 Hz. Before recording began, after conductive gel application, all sensors’ electrode impedance was confirmed to be at least below 50 kΩ.

## Results

To illustrate the functionality, typical workflow, specific tools, and visualizations of the SleepEEGpy pipeline, we performed preprocessing and analysis of overnight sleep EEG data obtained in 44 young, healthy adults participating in a memory consolidation study (Methods). Similar applications can be used for any type of sleep recording, such as a daytime nap or a study in clinical populations. In this study, each dataset consisted of a ∼7h undisturbed sleep hd-EEG (256-channel, EGI) combined with polysomnography including EOG, EMG, and video.

### Section A Preprocessing: cleaning and optional ICA

The first step in the pipeline is to perform cleaning (fig. 2a) via downsampling, filtering, visual annotation and interpolation of noisy electrodes, and identification and exclusion of noisy temporal intervals from subsequent analysis. We illustrate this on one representative dataset (27-year-old female). For the preprocessing we down-sampled the data to 250 Hz; and used common average reference. Additionally, we choose to perform ICA (see below). In terms of filtering, we used a band-pass filter of 0.75–40 Hz, without applying a specific notch filter.

#### Code snippet #1: A minimal working example (MWE) of the cleaning tool in the pipeline

Import the tool (line 1), initialize it with paths to the EEG signal and output folder (l. 2-4), resample and filter the signal (l. 5-6), plot the signal to annotate bad channels and bad epochs (l. 7), and interpolate bad channels (l. 8).

~~~
1    from sleepeegpy.pipeline import CleaningPipe
2    pipe = CleaningPipe(
3        path_to_eeg=“./data/EL3001_SLEEP.mff”,
4        output_dir=“./processed/EL3001”)
5    pipe.resample(sfreq=250)
6    pipe.filter(l_freq=0.75, h_freq=40)
7    pipe.plot(butterfly=True)
8    pipe.interpolate_bads(reset_bads=True)
~~~

Next, as part of the SleepEEGpy pipeline, we used an MNE-based data browser to annotate bad channels and bad temporal intervals. Manual inspection and annotation of raw signals were performed in the “butterfly” view and complemented by PSD topography inspection to finalize the exclusion of abnormal channels. In our example participant, 8.95% of the channels were annotated as bad and interpolated using MNE-based spherical spline interpolation (60). Across the entire dataset (N=44), 14.0 ± 4.6 % (mean ± STD) were marked as “bad channels”, and four subjects with more than 25% bad channels were excluded, leaving N=40 for subsequent analysis. Next, we marked temporal intervals in the representative dataset as “bad” (4.84% of the data in the example, 5.3 ± 2.4 % across the entire N=40 dataset). Next, sleep scoring was performed manually according to established American Academy of Sleep Medicine (AASM) criteria (3) to create a specific vector text file that could be fed to SleepEEGpy as input for subsequent preprocessing and analysis (Methods). The other 39 datasets were sleep scored automatically using YASA’s algorithm (41).

#### Code snippet #2: An MWE of the ICA tool in the pipeline

Import the tool (line 1), initialize it with the preceding object in the pipeline and the number of PCA components (l.2), run the ICA decomposition (l.3), plot the sources to annotate components for exclusion (l.4), apply the exclusion (l.5), and save the ICA decomposition and the cleaned EEG signal (l.6-7).

~~~
1    from sleepeegpy.pipeline import ICAPipe
2    ica_pipe = ICAPipe(prec_pipe=pipe, n_components=40)
3    ica_pipe.fit()
4    ica_pipe.plot_sources()
5    ica_pipe.apply()
6    ica_pipe.save_ica(“exclude-ica.fif”)
7    ica_pipe.save_raw(“cleaned_ica_raw.fif”)
~~~

Lastly, we applied ICA to identify and remove components of the EEG unrelated to brain activity. We chose to annotate and remove components associated with the electrocardiograph/heartbeat before proceeding to the analysis.

Figure 3 presents the dashboard, a visual summary after preprocessing. Here we used the representative dataset of participant EL3001 and plotted the activity at the Pz electrode. We created the dashboard to be a helpful, standardized visualization to evaluate the quality of the applied preprocessing. It includes the following subplots: general preprocessing information (figure 3a), topography of selected frequency ranges (3b), TFR with hypnogram before (3c) and after (3d) rejection of bad intervals and interpolation of bad electrodes, PSD plots of vigilance state before (3e) and after (3f) cleaning.

Because the cleaning (no ICA was applied) was successful, the topographic PSD maps of the dominant EEG rhythms of each vigilance state (figure 3b) show A) Wakefulness (top left) is characterized by maximal alpha (8-12 Hz) activity over the occipital lobe. B) In N2 sleep (top right), sigma (12-15 Hz) activity predominates over the centroparietal electrodes. C) In N3 sleep (bottom left), slow wave activity (<4 Hz) is maximal over the frontal cortex, and D) REM sleep (bottom right) is characterized by theta (4-8 Hz) activity with its signature scalp topography.

Furthermore, the hypnogram (fig. 3c and 3d) demonstrates a “typical” time-course of sleep/wake states. As expected, most bad and rejected temporal intervals are associated with wake intervals where locomotion and artifacts occur more readily (fig. 3d, marked as vertical white bars). The PSD after the preprocessing (fig 3f), compared to before (fig. 3e) shows expected signatures of vigilance states. Specifically, the slow-wave activity (SWA, power < 4Hz) is maximal in N3 sleep, lower in N2 sleep, lower in REM sleep, and lowest in wakefulness; showing that sigma (spindle) activity (12-15 Hz) is dominant in N2 and N3 sleep; highlighting alpha (8-12 Hz) peak in wakefulness, and diffuse theta (4-8 Hz) activity in REM sleep. Note that some of these features do not appear clearly before preprocessing (fig. 3e, left).

Thus, by comparing spectrograms and PSDs before and after cleaning, the user can effectively form an initial impression of the data quality and the effectiveness of the cleaning process (and whether additional iterations may be needed). Overall, the “dashboard” provides a visual summary of a specific dataset, its cleaning/preprocessing, and markers attesting to its quality, which constitute a useful first step for the investigator before proceeding to detailed analysis.

### Section B, analysis: the event-based and spectral-based tools

#### B1: Sleep spindle detection

To illustrate the event analysis tools (B1), we applied the YASA-based spindle detection in the N2 sleep of our representative dataset.

##### Code snippet #4: An MWE of the spindle detection and analysis tool in the pipeline

Import the tool (line 1), initialize it with the preceding object in the pipeline and the hypnogram file with a scoring frequency 1 Hz (l.2-5), and then detect spindles only in the N2 sleep stage (l.6). Plot the average spindle (l.7) and plot the topography distribution of the spindles’ amplitude, frequency, and duration (l.8-9). Next, compute the average TFR of the detected spindles per channel (l. 10-14), and finally plot it for the Pz (E101) channel (l. 15).

~~~
1    from sleepeegpy.pipeline import SpindlesPipe
2    spindles_pipe = SpindlesPipe(
3        prec_pipe=ica_pipe,
4        path_to_hypno=“./EL3001/hypno.txt”,
5        hypno_freq=1)
6    spindles_pipe.detect(include=(2))
7    pipe.results.plot_average()
8    spindles_pipe.plot_topomap_collage(
9        props=[“Amplitude”, “Frequency”, “Duration”])
10    spindles_pipe.compute_tfr(
11        freqs=(10, 20),
12        n_freqs=100,
13        time_before=1,
14        time_after=1)
15    spindles_pipe.tfrs[“N2”].plot([“E101”])
~~~

To detect spindle events, we used the default parameters of YASA. These included a 12-15 Hz spindle frequency range, 1-30 Hz broad band range, spindle duration between 0.5–2 s, and 500 ms as the minimal time interval for detecting two distinct spindles. The detection thresholds for a single spindle event were 0.2 relative power, 0.65 moving correlation, and 1.5 STDs above the mean of a moving root-mean-square of the sigma-filtered signal. The signal was re-referenced to a common average reference. With these default parameters, we detected 48,057 spindles across all EEG channels, corresponding to an average of ∼187 spindles per channel. Given that the duration of N2 sleep in this dataset was 193 minutes, this reflects a detection rate of ∼0.97 spindles/minute. This relatively low rate is reasonable given that all 257 channels were included. Since spindles are mostly detected over midline scalp electrodes and some electrodes, such as lateral or facial electrodes, only have marginal detections, their inclusion is bound to lower the average rate. The average time course of the detected spindles aligned at the peak is shown in Figure 4a, and Figure 4b depicts the average time–frequency decomposition (spectrogram) representation of the Pz (E101) channel, showing a slight decrease in spindle frequency from beginning to end, in line with previous findings (51). Figure 4c shows the topographical signatures of different spindle characteristics (left, amplitude; middle, frequency; right, duration), revealing established phenomena such as the prevalence of slower (<13Hz) spindles in frontal electrodes vs. fast (>13 Hz) spindles over centroparietal electrodes (61,62).

#### B2: Spectral tool

We performed MNE-based spectral analysis for the entire dataset separately for each sleep stage and frequency band (slow/delta, theta, alpha, sigma, beta; see Methods). First, we reviewed and edited the default MNE parameters to set FFT and hamming window length to 2048 samples and window overlap to 1024 samples.

##### Code snippet #5: An MWE of the grand spectral tool in the pipeline

First, import both the spectral and grand spectral tools (line 1-2). Initialize a spectral tool object for each subject (l.5-9). With these, initialize the grand spectral tool (l. 10). Next, compute PSD per sleep stage using the average reference, 2048 samples length of the FFT, and the Hamming window with an overlap of 1024 samples (l. 11-17). Plot scalp topography distribution of PSDs per sleep stage and PSD plot (l. 18-24). Finally, we parametrize the PSDs and plot fits of periodic and aperiodic components for the N2 sleep stage and Pz (E101) channel (l. 25-27).

~~~
1     from pathlib import Path
2     from sleepeegpy.pipeline import SpectralPipe, GrandSpectralPipe
3     p = Path(“./data”)
4     folders = [p/subj_num for subj_num in codes]
5     pipes = [SpectralPipe(
6               path_to_eeg=folder/”ICAPipe”/”cleaned_ica_raw.fif”,
7               output_dir=folder,
8               path_to_hypno=folder/”hypno.txt”,
9               hypno_freq=1) for folder in folders]
10     grand_pipe = GrandSpectralPipe(pipes=pipes, output_dir=p)
11     grand_pipe.compute_psd(
12          sleep_stages={“Wake”: 0, “N1”: 1, “N2”: 2, “N3”: 3, “REM”: 4},
13          reference=“average”,
14          n_fft=2048,
15          n_per_seg=2048,
16          n_overlap=1024,
17          window=“hamming”)
18     grand_pipe.plot_topomap_collage(
19          bands={
20               “Delta”: (1, 4),
21               “Theta”: (4, 8),
22               “Alpha”: (8, 12.5),
23               “Sigma”: (12, 15),
24               “Beta”: (12.5, 30)})
25     grand_pipe.plot_psds()
26     grand_pipe.parametrize(picks=[“E101”])
27     grand_pipe.fooofs[‘N2’].get_fooof(ind=0).plot(plot_peaks=‘dot’)
~~~

The resulting topographical distributions of PSDs per frequency band per sleep stage averaged over all 40 subjects are shown in Figure 5. In addition to typical activity signatures (described in fig. 3b, above), additional data features can be viewed and assessed here. For example, high-frequency beta activity is maximal during wakefulness; by contrast, delta activity during wakefulness shows hotspots around orbital electrodes, due to saccades “injecting” power into this frequency range.

Finally, we display the average PSD plot as a function of the sleep stage across the entire dataset (N=40) and apply parameterization to the PSDs (fig. 6). As observed in the representative dataset shown in the “dashboard” (fig. 3f), this plot reveals expected signatures of vigilance states such as SWA gradient N3>N2, diffuse theta activity in N1 and REM sleep, alpha activity in wakefulness, and maximal high-frequency (>20 Hz) activity in wakefulness. In addition, a peak in sigma activity was observed in the average PSD in N2/N3 sleep. This can also be demonstrated by identifying the periodic component’s peak frequency (∼13.5 Hz) using SpecParam (fig. 6b, rightmost vertical green line).

## Discussion

SleepEEGpy is an accessible and user-friendly tool for beginners in sleep EEG data analysis. It offers a comprehensive and user-friendly solution to support sleep EEG research from start to end by providing tools that enable preprocessing, analysis, and visualization of sleep EEG data. By leveraging the MNE-Python library and incorporating features from YASA and SpecParam (formerly FOOOF), it combines the advantages of general-purpose tools with those of specialized tools. Researchers can benefit from various functionalities, including artifact removal, ICA, event detection, and spectral analyses.

Developed as a streamlined introduction, it facilitates the learning curve for students and newcomers by unifying essential functionalities. New users should be able to run a standard pipeline without having to worry about the compatibility of data structures, coding errors and arbitrary parameter definitions. Therefore, they can focus on understanding the general high-level steps of analyzing sleep data. Furthermore, SleepEEGpy can be used for learning and teaching specific steps. For example, the dashboard’s standardized visualization makes it easy and fast to assess the quality of the preprocessing. Hence it can function as a useful tool to learn and improve manual preprocessing steps such as annotation of bad epochs and bad electrodes.

### Applying SleepEEGpy to overnight sleep data

We demonstrated a typical workflow of sleepEEGpy with continuous hd-EEG data from healthy young. With the dashboard (fig. 3), we summarized and assessed the quality of the preprocessing. The minimal preprocessing included resampling, bandpass filtering, electrode interpolation and epoch rejection. Finally, an ICA was applied to regress out the components related to electrocardiography/heartbeat. An examination of the most important oscillatory patterns that typically occur during sleep indicates that the sleep data has been successfully pre-processed. We further illustrated that with SleepEEGpy it is possible to reliably analyze events during sleep by detecting sleep spindles (fig. 4). We presented group-averages of the topographical distribution of PSD per frequency band and sleep stage (fig. 5). And finally, we separately inspect the average PSD for each vigilance state, averaged across our participant cohort (fig. 6). In summary, SleepEEGpy provides an easy way to perform general analysis and visualization of raw sleep EEG data up to overviews of group averages.

### Comparison with other software packages and advantages

of SleepEEGpy. The main advantages of SleepEEGpy relative to other available software tools are its simplicity and all-in-one functionality. At present, EEG software packages are either implemented in MATLAB and behind a paywall (26,27,29,30,34,35,38), are optimized for sleep EEG but restricted to either preprocessing (45), sleep scoring (64), or specific analyses (41), or are based on Python but not necessarily optimized for sleep research (28). Thus, SleepEEGpy helps to address an unmet need by providing a comprehensive package that goes beyond the typical configuration in many labs that combine multiple software environments to work with sleep EEG data. Its free open-source nature ensures accessibility to students and sleep research labs.

Specifically, if we compare SleepEEGpy to the software packages it is based on, SleepEEGpy’s advantages include start-to-end in one pipeline and pre-made yet customizable visualizations. It offers a simplified yet comprehensive framework for easy applicable sleep data analysis for inexperienced users. Nevertheless, most of YASA and MNE’s meta-parameters (e.g. filter type or spindle detection threshold) can be passed to the underlying function through the SleepEEGpy framework. Naturally, with time the initially inexperienced user might out-grow SleepEEGpy and at a certain level of expertise work directly with the specialized toolboxes and their extended functionality.

### Ease-of-use

SleepEEGpy requires only basic knowledge of Python syntax and Jupyter notebooks. The pipeline is based mostly on classes and their methods, and the most complicated task an average user can encounter is writing a ‘for’ loop to optimize their pipeline. Jupyter notebooks are implemented for each stage of the pipeline so that running the tools is nearly automatic and includes embedded explanations at each step. The code repository and notebooks can be found at https://github.com/NirLab-TAU/sleepeegpy. SleepEEGpy is accompanied with a webpage providing detailed API documentation and published notebooks. The documentation is built using Sphinx (65) and hosted on GitHub. The example dataset can be found at Zenodo (66): 10.5281/zenodo.10362190. An end-to-end example of dataset retrieval, preprocessing, and analysis can be found in the “complete pipeline” notebook.

### Limitations and future directions

SleepEEGpy could be improved by including additional analysis methods, for example a module for statistics (parametric and non-parametric tests) or a module for source estimation. Data type compatibility of SleepEEGpy could be further extended, for example with the EEG-BIDS data structure (67). We believe that the code availability and free software licenses (SleepEEGpy is released under the MIT license) allow rapid expansion of such functionalities by the community. An additional limitation is the required knowledge of MNE and YASA for more advanced analysis methods. As of now, if the analysis prompts for an adjustment of a meta-parameter (e.g. spindle detection threshold or specific bandpass filtering), the details are best described in the documentation of MNE or YASA itself. However, for a new user, the defaults should be sufficient to get to know the processing of sleep EEG. All these limitations could, in principle, be addressed with additional code and documentation. Hence, we hope that this package will be further developed by the growing community of sleep investigators committed to open science and high-quality open-source software. With the increased use of Python as the preferred programming language and its interface with machine-learning tools, we envision SleepEEGpy as an ideal entry point to become familiar with sleep EEG analysis.

### Conclusion

Overall, SleepEEGpy simplifies the process of sleep EEG analysis by providing a unified and user-friendly interface for new users. It reduces the burden of learning and using multiple software packages, allowing researchers to focus more on their scientific questions and obtain valuable insights from sleep EEG data.

## Acknowledgments

We thank Gal Zatelman, May Eliyahu, and Shir Frank for their assistance with data collection and sleep scoring and Dr. Noa Bar-Ilan Regev for administrative assistance. This study was supported by ERC-2019-CoG 864353 and a grant from the Aufzien Family Center for the Prevention and Treatment of Parkinson’s Disease (Y.N.).

## Author contributions

Y.N. conceived the package development and secured funding. G.B. designed and programmed SleepEEGpy, with assistance from R.F. and F.J.S.; M.A., V.Z., and R.S. provided user experience insights. E.B. led the sleep and memory consolidation project (EEG datasets). G.B. performed data preprocessing and analysis. G.B, R.F., F.J.S., and Y.N. wrote the manuscript. All authors provided feedback, assisted with testing the tool during development, and commented on the manuscript.

## Notes

https://zenodo.org/records/10362190

### Competing Interest Statement

The authors have declared no competing interest.

### Summary of Updates

We improved the quality of the figure (the conversion from word was blurry).

## References

1. Kryger MH, Roth T. Principles and Practice of Sleep Medicine. 6. edition. Philadelphia, PA: Elsevier - Health Sciences Division (2016). 1784 p.

2. Nir Y, de Lecea L. Sleep and vigilance states: Embracing spatiotemporal dynamics. Neuron (2023) 111:1998–2011. doi: 10.1016/j.neuron.2023.04.012

3. Berry RB, Brooks R, Gamaldo C, Harding SM, Lloyd RM, Quan SF, Troester MT, Vaughn BV. AASM Scoring Manual Updates for 2017 (Version 2.4). Journal of Clinical Sleep Medicine (2017) 13:665–666. doi: 10.5664/jcsm.6576

4. Gaiduk M, Serrano Alarcón Á, Seepold R, Martínez Madrid N. Current status and prospects of automatic sleep stages scoring: Review. Biomed Eng Lett (2023) 13:247–272. doi: 10.1007/s13534-023-00299-3

5. Urtnasan E, Park J-U, Joo EY, Lee K-J. Deep Convolutional Recurrent Model for Automatic Scoring Sleep Stages Based on Single-Lead ECG Signal. Diagnostics (Basel) (2022) 12:1235. doi: 10.3390/diagnostics12051235

6. Fiorillo L, Puiatti A, Papandrea M, Ratti P-L, Favaro P, Roth C, Bargiotas P, Bassetti CL, Faraci FD. Automated sleep scoring: A review of the latest approaches. Sleep Medicine Reviews (2019) 48:101204. doi: 10.1016/j.smrv.2019.07.007

7. Cox R, Fell J. Analyzing human sleep EEG: A methodological primer with code implementation. Sleep Medicine Reviews (2020) 54:101353. doi: 10.1016/j.smrv.2020.101353

8. Geva-Sagiv M, Nir Y. Local Sleep Oscillations: Implications for Memory Consolidation. Front Neurosci (2019) 13:813. doi: 10.3389/fnins.2019.00813

9. Geva-Sagiv M, Mankin EA, Eliashiv D, Epstein S, Cherry N, Kalender G, Tchemodanov N, Nir Y, Fried I. Augmenting hippocampal–prefrontal neuronal synchrony during sleep enhances memory consolidation in humans. Nat Neurosci (2023) 26:1100–1110. doi: 10.1038/s41593-023-01324-5

10. Schmidig JF, Geva-Sagiv M, Falach R, Yakim S, Gat Y, Sharon O, Fried I, Nir Y. A visual paired associate learning (vPAL) paradigm to study memory consolidation during sleep. [preprint]. Neuroscience (2023). doi: 10.1101/2023.03.28.534494

11. Gais S, Mölle M, Helms K, Born J. Learning-Dependent Increases in Sleep Spindle Density. J Neurosci (2002) 22:6830–6834. doi: 10.1523/JNEUROSCI.22-15-06830.2002

12. Marshall L, Born J. The contribution of sleep to hippocampus-dependent memory consolidation. Trends in Cognitive Sciences (2007) 11:442–450. doi: 10.1016/j.tics.2007.09.001

13. Schabus M, Gruber G, Parapatics S, Sauter C, Klösch G, Anderer P, Klimesch W, Saletu B, Zeitlhofer J. Sleep Spindles and Their Significance for Declarative Memory Consolidation. Sleep (2004) 27:1479–1485. doi: 10.1093/sleep/27.7.1479

14. Kurth S, Ringli M, Geiger A, LeBourgeois M, Jenni OG, Huber R. Mapping of Cortical Activity in the First Two Decades of Life: A High-Density Sleep Electroencephalogram Study. J Neurosci (2010) 30:13211–13219. doi: 10.1523/JNEUROSCI.2532-10.2010

15. Helfrich RF, Mander BA, Jagust WJ, Knight RT, Walker MP. Old Brains Come Uncoupled in Sleep: Slow Wave-Spindle Synchrony, Brain Atrophy, and Forgetting. Neuron (2018) 97:221-230.e4. doi: 10.1016/j.neuron.2017.11.020

16. Siclari F, Baird B, Perogamvros L, Bernardi G, LaRocque JJ, Riedner B, Boly M, Postle BR, Tononi G. The neural correlates of dreaming. Nat Neurosci (2017) 20:872–878. doi: 10.1038/nn.4545

17. Mander BA, Marks SM, Vogel JW, Rao V, Lu B, Saletin JM, Ancoli-Israel S, Jagust WJ, Walker MP. β-amyloid disrupts human NREM slow waves and related hippocampus-dependent memory consolidation. Nat Neurosci (2015) 18:1051–1057. doi: 10.1038/nn.4035

18. Ferrarelli F. Sleep spindles as neurophysiological biomarkers of schizophrenia. Eur J of Neuroscience (2023) doi: 10.1111/ejn.16178

19. Tucker DM, Waters AC, Holmes MD. Transition from cortical slow oscillations of sleep to spike-wave seizures. Clinical Neurophysiology (2009) 120:2055–2062. doi: 10.1016/j.clinph.2009.07.047

20. Kaya İ. “A Brief Summary of EEG Artifact Handling.,” Brain-Computer Interface. IntechOpen (2021) doi: 10.5772/intechopen.99127

21. Cohen MX. Analyzing Neural Time Series Data. MIT Press (2014). 600 p. doi: 10.7551/mitpress/9609.001.0001

22. Delorme A. EEG is better left alone. Sci Rep (2023) 13:2372. doi: 10.1038/s41598-023-27528-0

23. Riedner BA, Vyazovskiy VV, Huber R, Massimini M, Esser S, Murphy M, Tononi G. Sleep Homeostasis and Cortical Synchronization: III. A High-Density EEG Study of Sleep Slow Waves in Humans. Sleep (2007) 30:1643–1657. doi: 10.1093/sleep/30.12.1643

24. Das RK, Martin A, Zurales T, Dowling D, Khan A. A Survey on EEG Data Analysis Software. Sci (2023) 5:23. doi: 10.3390/sci5020023

25. Tadel F, Baillet S, Mosher JC, Pantazis D, Leahy RM. Brainstorm: A User-Friendly Application for MEG/EEG Analysis. Computational Intelligence and Neuroscience (2011) 2011:e879716. doi: 10.1155/2011/879716

26. Delorme A, Makeig S. EEGLAB: an open source toolbox for analysis of single-trial EEG dynamics including independent component analysis. Journal of Neuroscience Methods (2004) 134:9–21. doi: 10.1016/j.jneumeth.2003.10.009

27. Oostenveld R, Fries P, Maris E, Schoffelen J-M. FieldTrip: Open Source Software for Advanced Analysis of MEG, EEG, and Invasive Electrophysiological Data. Computational Intelligence and Neuroscience (2010) 2011:e156869. doi: 10.1155/2011/156869

28. Gramfort A. MEG and EEG data analysis with MNE-Python. Front Neurosci (2013) 7: doi: 10.3389/fnins.2013.00267

29. Nolan H, Whelan R, Reilly RB. FASTER: Fully Automated Statistical Thresholding for EEG artifact Rejection. Journal of Neuroscience Methods (2010) 192:152–162. doi: 10.1016/j.jneumeth.2010.07.015

30. Mognon A, Jovicich J, Bruzzone L, Buiatti M. ADJUST: An automatic EEG artifact detector based on the joint use of spatial and temporal features. Psychophysiology (2011) 48:229–240. doi: 10.1111/j.1469-8986.2010.01061.x

31. Bigdely-Shamlo N, Mullen T, Kothe C, Su K-M, Robbins KA. The PREP pipeline: standardized preprocessing for large-scale EEG analysis. Frontiers in Neuroinformatics (2015) 9: doi: 10.3389/fninf.2015.00016

32. ‘t Wallant DC, Muto V, Gaggioni G, Jaspar M, Chellappa SL, Meyer C, Vandewalle G, Maquet P, Phillips C. Automatic artifacts and arousals detection in whole-night sleep EEG recordings. Journal of Neuroscience Methods (2016) 258:124–133. doi: 10.1016/j.jneumeth.2015.11.005

33. Jas M, Engemann DA, Bekhti Y, Raimondo F, Gramfort A. Autoreject: Automated artifact rejection for MEG and EEG data. NeuroImage (2017) 159:417–429. doi: 10.1016/j.neuroimage.2017.06.030

34. da Cruz JR, Chicherov V, Herzog MH, Figueiredo P. An automatic pre-processing pipeline for EEG analysis (APP) based on robust statistics. Clinical Neurophysiology (2018) 129:1427–1437. doi: 10.1016/j.clinph.2018.04.600

35. Gabard-Durnam LJ, Mendez Leal AS, Wilkinson CL, Levin AR. The Harvard Automated Processing Pipeline for Electroencephalography (HAPPE): Standardized Processing Software for Developmental and High-Artifact Data. Front Neurosci (2018) 12: doi: 10.3389/fnins.2018.00097

36. Bailey NW, Biabani M, Hill AT, Miljevic A, Rogasch NC, McQueen B, Murphy OW, Fitzgerald PB. Introducing RELAX: An automated pre-processing pipeline for cleaning EEG data - Part 1: Algorithm and application to oscillations. Clinical Neurophysiology (2023) 149:178–201. doi: 10.1016/j.clinph.2023.01.017

37. Somervail R, Cataldi J, Stephan AM, Siclari F, Iannetti GD. Dusk2Dawn: an EEGLAB plugin for automatic cleaning of whole-night sleep electroencephalogram using Artifact Subspace Reconstruction. Sleep (2023) doi: 10.1093/sleep/zsad208

38. Blum S, Jacobsen NSJ, Bleichner MG, Debener S. A Riemannian Modification of Artifact Subspace Reconstruction for EEG Artifact Handling. Front Hum Neurosci (2019) 13: doi: 10.3389/fnhum.2019.00141

39. Anderer P, Roberts S, Schlögl A, Gruber G, Klösch G, Herrmann W, Rappelsberger P, Filz O, Barbanoj MJ, Dorffner G, et al. Artifact Processing in Computerized Analysis of Sleep EEG – A Review. Neuropsychobiology (1999) 40:150–157. doi: 10.1159/000026613

40. Desjardins JA, van Noordt S, Huberty S, Segalowitz SJ, Elsabbagh M. EEG Integrated Platform Lossless (EEG-IP-L) pre-processing pipeline for objective signal quality assessment incorporating data annotation and blind source separation. Journal of Neuroscience Methods (2021) 347:108961. doi: 10.1016/j.jneumeth.2020.108961

41. Vallat R, Walker MP. An open-source, high-performance tool for automated sleep staging. eLife (2021) 10:e70092. doi: 10.7554/eLife.70092

42. Donoghue T, Haller M, Peterson EJ, Varma P, Sebastian P, Gao R, Noto T, Lara AH, Wallis JD, Knight RT, et al. Parameterizing neural power spectra into periodic and aperiodic components. Nat Neurosci (2020) 23:1655–1665. doi: 10.1038/s41593-020-00744-x

43. Kluyver T, Ragan-Kelley B, Pérez F, Granger B, Bussonnier M, Frederic J, Kelley K, Hamrick J, Grout J, Corlay S, et al. Jupyter Notebooks – a publishing format for reproducible computational workflows. In: Loizides F, Scmidt B, editors. IOS Press (2016). p. 87–90 doi: 10.3233/978-1-61499-649-1-87

44. Andrillon T, Nir Y, Cirelli C, Tononi G, Fried I. Single-neuron activity and eye movements during human REM sleep and awake vision. Nat Commun (2015) 6:7884. doi: 10.1038/ncomms8884

45. Leach S, Sousouri G, Huber R. ‘High-Density-SleepCleaner’: An open-source, semiautomatic artifact removal routine tailored to high-density sleep EEG. Journal of Neuroscience Methods (2023) 391:109849. doi: 10.1016/j.jneumeth.2023.109849

46. Hyvarinen A. Fast and robust fixed-point algorithms for independent component analysis. IEEE Trans Neural Netw (1999) 10:626–634. doi: 10.1109/72.761722

47. Lee T-W, Girolami M, Sejnowski TJ. Independent Component Analysis Using an Extended Infomax Algorithm for Mixed Subgaussian and Supergaussian Sources. Neural Computation (1999) 11:417–441. doi: 10.1162/089976699300016719

48. Ablin P, Cardoso J-F, Gramfort A. Faster Independent Component Analysis by Preconditioning With Hessian Approximations. IEEE Trans Signal Process (2018) 66:4040–4049. doi: 10.1109/TSP.2018.2844203

49. Hansson-Sandsten M. Optimal Multitaper Wigner Spectrum Estimation of a Class of Locally Stationary Processes Using Hermite Functions. EURASIP J Adv Signal Process (2010) 2011:1–15. doi: 10.1155/2011/980805

50. Lacourse K, Delfrate J, Beaudry J, Peppard P, Warby SC. A sleep spindle detection algorithm that emulates human expert spindle scoring. Journal of Neuroscience Methods (2019) 316:3–11. doi: 10.1016/j.jneumeth.2018.08.014

51. Andrillon T, Nir Y, Staba RJ, Ferrarelli F, Cirelli C, Tononi G, Fried I. Sleep Spindles in Humans: Insights from Intracranial EEG and Unit Recordings. J Neurosci (2011) 31:17821–17834. doi: 10.1523/JNEUROSCI.2604-11.2011

52. Purcell SM, Manoach DS, Demanuele C, Cade BE, Mariani S, Cox R, Panagiotaropoulou G, Saxena R, Pan JQ, Smoller JW, et al. Characterizing sleep spindles in 11,630 individuals from the National Sleep Research Resource. Nat Commun (2017) 8:15930. doi: 10.1038/ncomms15930

53. The pandas development team. pandas-dev/pandas: Pandas. (2023) doi: 10.5281/ZENODO.3509134

54. Massimini M, Huber R, Ferrarelli F, Hill S, Tononi G. The Sleep Slow Oscillation as a Traveling Wave. J Neurosci (2004) 24:6862–6870. doi: 10.1523/JNEUROSCI.1318-04.2004

55. Nir Y, Staba RJ, Andrillon T, Vyazovskiy VV, Cirelli C, Fried I, Tononi G. Regional Slow Waves and Spindles in Human Sleep. Neuron (2011) 70:153–169. doi: 10.1016/j.neuron.2011.02.043

56. Agarwal R, Takeuchi T, Laroche S, Gotman J. Detection of Rapid-Eye Movements in Sleep Studies. IEEE Trans Biomed Eng (2005) 52:1390–1396. doi: 10.1109/TBME.2005.851512

57. Tallon-Baudry C, Bertrand O, Delpuech C, Pernier J. Oscillatory γ-Band (30–70 Hz) Activity Induced by a Visual Search Task in Humans. J Neurosci (1997) 17:722–734. doi: 10.1523/JNEUROSCI.17-02-00722.1997

58. Slepian D. Prolate Spheroidal Wave Functions, Fourier Analysis, and Uncertainty-V: The Discrete Case. Bell System Technical Journal (1978) 57:1371–1430. doi: 10.1002/j.1538-7305.1978.tb02104.x

59. Welch P. The use of fast Fourier transform for the estimation of power spectra: A method based on time averaging over short, modified periodograms. IEEE Trans Audio Electroacoust (1967) 15:70–73. doi: 10.1109/TAU.1967.1161901

60. Perrin F, Pernier J, Bertrand O, Echallier JF. Spherical splines for scalp potential and current density mapping. Electroencephalography and Clinical Neurophysiology (1989) 72:184–187. doi: 10.1016/0013-4694(89)90180-6

61. Cox R, Schapiro AC, Manoach DS, Stickgold R. Individual Differences in Frequency and Topography of Slow and Fast Sleep Spindles. Front Hum Neurosci (2017) 11: doi: 10.3389/fnhum.2017.00433

62. Mölle M, Bergmann TO, Marshall L, Born J. Fast and Slow Spindles during the Sleep Slow Oscillation: Disparate Coalescence and Engagement in Memory Processing. Sleep (2011) 34:1411–1421. doi: 10.5665/SLEEP.1290

63. Ferrarelli F, Huber R, Peterson MJ, Massimini M, Murphy M, Riedner BA, Watson A, Bria P, Tononi G. Reduced Sleep Spindle Activity in Schizophrenia Patients. AJP (2007) 164:483–492. doi: 10.1176/ajp.2007.164.3.483

64. Combrisson E, Vallat R, Eichenlaub J-B, O’Reilly C, Lajnef T, Guillot A, Ruby PM, Jerbi K. Sleep: An Open-Source Python Software for Visualization, Analysis, and Staging of Sleep Data. Front Neuroinform (2017) 11: doi: 10.3389/fninf.2017.00060

65. Komiya T, Brandl G, Jean-François B, Shimizukawa T, Turner A, Andersen JL, Neuhäuser D, Finucane S, Lehmann R, Kampik T, et al. sphinx-doc/sphinx: Sphinx 7.2.6. (2023) doi: 10.5281/ZENODO.7857310

66. European Organization For Nuclear Research, OpenAIRE. Zenodo: Research. Shared. (2013) doi: 10.25495/7GXK-RD71

67. Pernet CR, Appelhoff S, Gorgolewski KJ, Flandin G, Phillips C, Delorme A, Oostenveld R. EEG-BIDS, an extension to the brain imaging data structure for electroencephalography. Sci Data (2019) 6:103. doi: 10.1038/s41597-019-0104-8

